# Integrative taxonomy of *Trichiurus* (Scombriformes: Trichiuridae) reveals a new cutlassfish species from Java, Indonesia

**DOI:** 10.64898/2026.05.05.722933

**Authors:** Tianqin Wu, Chenhong Li

## Abstract

The genus *Trichiurus* is the most economically valuable fish in the family Trichiuridae, currently recognized to include 10 valid species. However, historically numerous morphologically similar congeners have been erroneously assigned as synonyms or subspecies of *T. lepturus*. In this study, we examined 16 hairtail specimens collected from the southern waters of Java Island, Indonesia. Integrated morphological and mitochondrial phylogenetic analyses (COX1 and 16S rRNA), compared against global *Trichiurus* sequences, revealed that these specimens form an independent lineage that diverged early from other congeners. Consequently, we describe this lineage as a previously undescribed cryptic species. Diagnostic characters include: first anal-fin spine below 36th–37th dorsal-fin rays; anus below 35th–36th dorsal-fin rays; anteriormost tip of supraoccipital well posterior to posterior distal margin of eye; anterior margin of the pectoral-fin spine non-serrated; fangs on both jaws with barb-like processes; upper jaw long, mean 16.6% (15.5–17.6%) of preanal length; snout short, 12.0% (10.9–13.1%) of preanal length; eye small, diameter 5.3% (4.3–5.7%) of preanal length; and absence of hyperostosis on dorsal cranium. We herein propose the name ***Trichiurus javaensis* sp. nov**., and provide a formal morphological description and diagnostic characterization of this species.

## Introduction

The taxonomy of the cutlassfishes (Trichiuridae, Rafinesque, 1810) has a long and complicated history dating back more than 250 years, when *Trichiurus lepturus* (Linnaeus, 1758) was first discribed. Under the taxonomic framework of Nakamura and Parin (1993), the family is divided into nine genera. Five genera (*Aphanopus*, *Assurger*, *Benthodesmus*, *Evoxymetopon*, and *Lepidopus*) are distinguished by forked caudal fins, while four others (*Lepturacanthus*, *Tentoriceps*, *Eupleurogrammus*, and *Trichiurus*), as well as the more recently established genus *Demissolinea* (Burhanuddin & Iwatsuki, 2003), are characterized by reduced or vestigial caudal fin structures. Despite being the most geographically widespread and commercially exploited genus, *Trichiurus* reminas systematically problematic due to morphological ambiguities and unresolved species complexes. This lack of clarity impedes effective fisheries management, biodiversity assessment, and species identification in trade.

*Trichiurus lepturus* was described by Linnaeus (1758) when taxonomic understanding of this group was severely limited. Based on the re-examination and morphological documentation of Linnaeus’ type material by Li (2006), the four syntype specimens deposited in the Swedish Museum of Natural History (NRM 15) actually belong to the genus *Lepturacanthus*, while specimen ZIU 172 held at Uppsala University is assignable to the genus *Eupleurogrammus*. Only the two specimens from the Atlantic United States and Jamaica correspond to *T. lepturus* as currently circumscribed.

Over the following century, numerous putative new species were described. Prior to the 20th century, due to the difficulty of conducting comparative morphological analyses on morphologically conserved cutlassfish taxa, numerous nominal species described were subsequently relegated to junior synonyms of *T. lepturus*, as their original descriptions failed to provide diagnostic characters sufficient to reliably distinguish them from the nominotypical species. *Clupea haumela* Forsskål, 1775 described from the Red Sea is currently widely accepted as a junior synonym of *T. lepturus*. However, pre-20th century studies recognized *T. lepturus* as the Atlantic cutlassfish and *T. haumela* as the corresponding species from the Indian Ocean and Indo-West Pacific. This taxonomic treatment was not supported by substantive evidence, but rather arising from the limited biogeographic and taxonomic information available at the time. *Trichiurus argenteus* Shaw, 1803 (Argentina) and *Trichiurus coxii* Ramsay & Ogilby, 1887, (New South Wales, Australia) are both treated as junior synonyms of *T. lepturus* based on their original morphological descriptions. *Trichiurus lajor* Bleeker, 1854 (Sulawesi, Indonesia) and *Trichiurus malabaricus* Day, 1865 (Cochin, India) were assigned as junior synonyms of *T. haumela* by de Beaufort (1951) and Day (1876), respectively (Tucker 1956). However, the original descriptions of both taxa documented the diagnostic trait of yellow coloration on the dorsal or pectoral fins, and the only valid species within the genus *Trichiurus* confirmed to possess yellow fins is *T. nanhaiensis*. Furthermore, *Trichiurus lepturus japonicus* Temminck & Schlegel, 1844 described based on specimens collected from Japan, was initially assigned only subspecific rank due to perceived morphological similarity to the nominotypical taxon. *Trichiurus savala* Cuvier, 1829 was subsequently assigned to *Lepturacanthus* Fowler, 1905, a subgenus originally erected under *Trichiurus*. Diagnostic characters of this subgenus include an elongated first anal-fin spine exceeding half the eye diameter and a slit on the lower jaw to accommodate the anteriorly projecting canine teeth of the upper jaw. However, following the taxonomic revision by Tucker (1956), *T. japonicus* was recognized as a distinct valid species rather than a subspecies, and *Lepturacanthus* was elevated to full generic rank instead of being retained as a subgenus. Nevertheless, these taxonomic treatments are arbitrary and not supported by robust, substantive evidence.

During the 20th century, taxonomic research on cutlassfishes yielded notable discrepancies between researchers from East Asia (China and Japan) and those from Europe, North America, and India, primarily driven by language barriers and restricted information exchange. In China, Chu (1931) proposed that only a single cutlassfish species, *T. haumela*, occurred along China’s coastal waters, whereas Liu contended that the populations belonged to *T. japonicus*. Tucker (1956) compared specimens of *T. haumela* from Shanghai and *T. japonicus* from Zhejiang Province, with *T. lepturus*; detecting no substantial morphological divergence, he concluded that there was only one valid cutlassfish species globally, *T. lepturus*, with all others treated as junior synonyms. This view was subsequently adopted by James (1965) from India, who asserted that cutlassfish populations along India’s coastal waters were conspecific with *T. lepturus*. *Trichiurus nanhaiensis* Wang & You, 1991 was described from the South China Sea. Diagnostic characters include a prominent knob-like bony protuberance on the dorsal cranium, yellow coloration of the dorsal fin, and a stouter, shorter caudal region. These traits enable reliable differentiation of this species from *T. japonicus* distributed in the Yellow Sea and East China Sea. However, as the original description was published in Chinese, this taxon is frequently misclassified as *T. lepturus nanhaiensis* (e.g., in the NCBI database). This misassignment stems from the fact that a sister lineage of *T. nanhaiensis* distributed in the Indian Ocean was identified as *T. lepturus* in studies by regional researchers (Yi et al., 2022), following Nakamura & Parin (1995).

Ten valid species of the genus *Trichiurus* are currently recognized worldwide. *Trichiurus auriga* and *T. gangeticus* are distinguished by unique features: *T. auriga* possesses fangs without barbs on both jaws, and *T. gangeticus* exhibits a serrated anterior margin on the pectoral-fin spine. The remaining eight species form two primary complexes based on shared morphology. The *T. lepturus* complex (*T. lepturus*, *T. japonicus*, *T. nanhaiensis*, *T. nitens*) is characterized by the anteriormost tip of the supraoccipital being situated well behind the posterior margin of the eye. In contrast, the *T. brevis* complex (*T. brevis*, *T. russelli*, *T. nickolensis*, *T. australis*) has the highest point of the supraoccipital positioned vertically above the eye’s posterior margin.

Key diagnostic feature include the vertical alignment of the anus or first anal-fin spine with specific dorsal-fin rays. For instance, the anus is located below the 34th–35th dorsal-fin ray in *T. brevis*, but below the 39th–40th in *T. lepturus*. *Trichiurus nitens* Garman, 1899, redescribed by Burhanuddin and Parin (2008), is a valid species distributed along the Pacific coasts of the Americas; its anus is positioned below the 35th–39th dorsal fin ray. However, these subtle and often overlapping morphological characters complicate reliable identification, underscoring the limitations of traditional morphology and the need for integrative taxonomic approaches.

The waters of Java Island, Indonesia, lie within the Coral Triangle, a global epicenter of marine biodiversity where species discoveries are common. However, the diversity of the cutlassfish genus *Trichiurus* in this region remains poorly studied. Specimens collected from these waters were morphologically consistent with the genus *Trichiurus* and generally similar to *T. lepturus*, the species to which they are commonly misidentified in fisheries. A detailed morphological examination revealed that these specimens possess a unique combination of diagnostic traits, such as the position of the anus and the eye diameter ratio, which are inconsistent with all currently described species.

Here, we describe these specimens as a new species based on integrated morphological and molecular evidence. We provide mitochondrial COX1 (cytochrome c oxidase subunit I) and 16s rRNA gene sequences to demonstrate their genetic distinctness from other recognized, morphologically similar trichiurid species. This work contributes to resolving the long-standing taxonomic confusion within the genus *Trichiurus*.

## Materials and Methods

Counts and measurements followed Hubbs and Lagler (1958). Measurements were taken from the left side of specimens using a digital caliper to an accuracy of 0.1 mm; meristic counts were also recorded. Because the caudal fin in hairtails is often damaged or lost, all body proportions are expressed as a percentage of preanal length instead of standard length.

The holotype was fixed in a 10% formalin solution for one month before being permanently preserved in 70% ethanol. Tissue samples (right pectoral fins) from additional specimens were preserved directly in 70% ethanol and stored at 4°C. Genomic DNA was extracted from these tissue samples using the Ezup Column Animal Genomic DNA Purification Kit (Sangon, Shanghai, China), following the manufacturer’s protocol.

The mitochondrial COX1 gene was amplified with the primers FishF1 (5′-CTACAACCCACCGCTTACTC-3′) and FishR1 (5′-GTTGTAATAAAGTTAATGGCGC-3′), and the 16S rRNA gene with the primers L2510 (5′-GCCTGTTTAACAAAAACAT-3′) and H3059 (5′-CGGTCTGAACTCAGATCACGT-3′) (Miya and Nishida, 1996). The PCR protocol consisted of an initial denaturation at 94°C for 5 min; 35 cycles of denaturation at 94°C for 30 s, annealing at 55°C for 45 s, and elongation at 72°C for 1 min; followed by a final elongation at 72°C for 5 min.

Successfully amplified products were purified and sequenced bidirectionally using Sanger sequencing at GENEWIZ (Shanghai, China). All available COX1 and 16S rRNA sequences for geographically verified species within the Trichiuridae family were downloaded from NCBI (accession numbers listed in Supplementary Table 1). A phylogenetic tree was reconstructed using MEGA software with the Maximum Likelihood method based on the General Time Reversible (GTR) model. Node support was assessed with 1000 bootstrap replicates, and a 50% site coverage cutoff was applied for gap/missing data treatment.

## Results

### Description

***Trichiurus javaensis* (Anonymous), 2025, sp. nov**

LSID urn:lsid:zoobank.org:act:E859DD81-EB05-4D5D-9035-4D2BD838777B

### Type materials

#### Holotype

SOU18011027-16 (Figure 1 A), adult female, 120.2 cm standard length (SL), 47.2 cm preanal length (PL); Indonesia: southern Indian Ocean, off the south coast of Java Island, Yogyakarta (8°10’S, 110°30’E), 5 February 2025, collected by the Anonymous aboard the fishing vessel KM. HARAPAN KITA 003994. Deposited in the Ichthyological Collection, Shanghai Ocean University (SOU), Shanghai, China. The holotype was fixed in 10% formalin solution and subsequently preserved in 75% ethanol for long-term storage.

**Figure 1.**
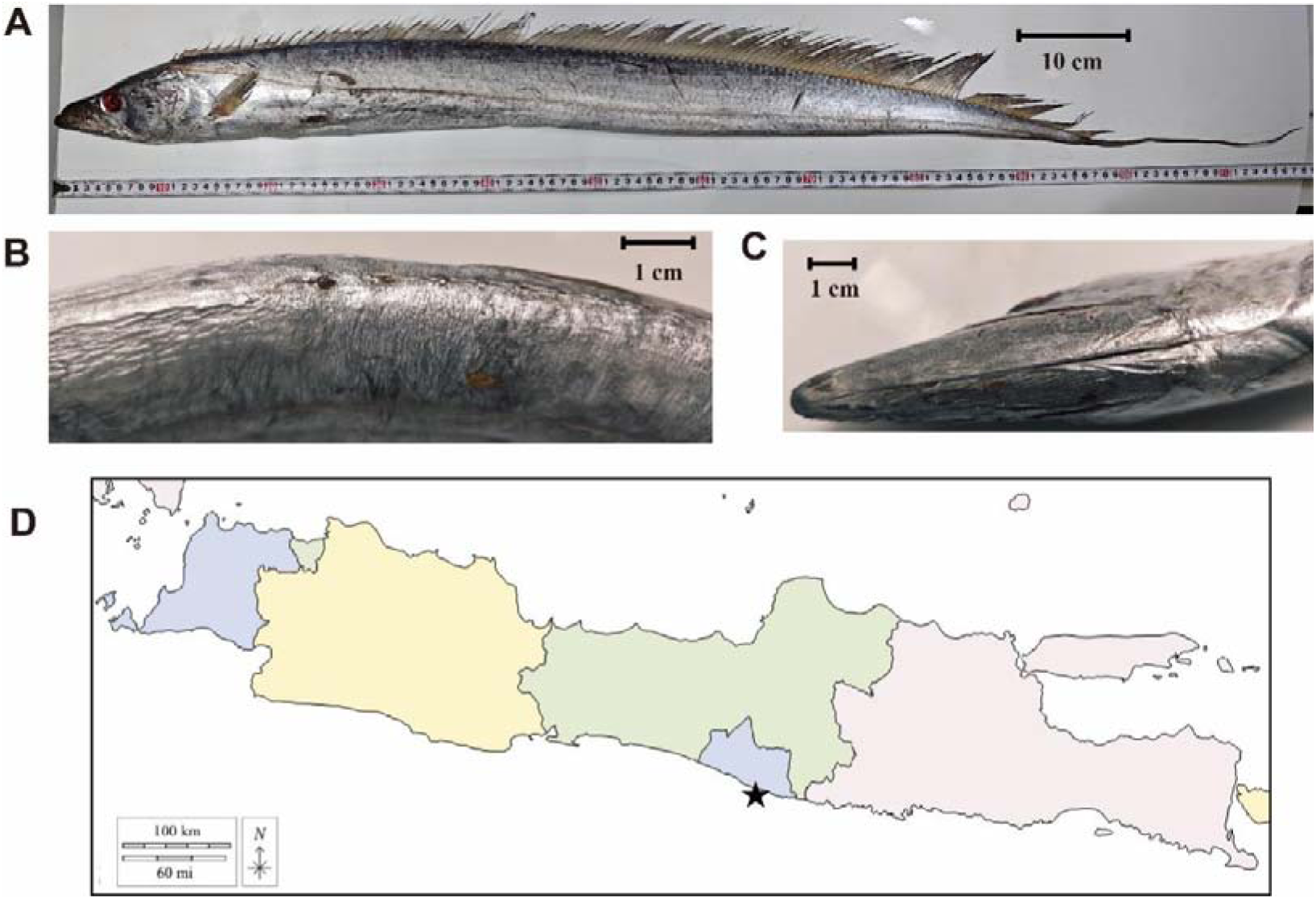
*Trichiurus javaensis*, holotype (SOU18011027-16). **A.** Lateral view. **B.** Close-up of the anus and first anal-fin spine. **C.** Ventral view of lower jaw, showing absence of slit. **D.** Type locality of new *Trichiurus* species located along southern coast of Java Island, Indonesia

**Figure 2.**
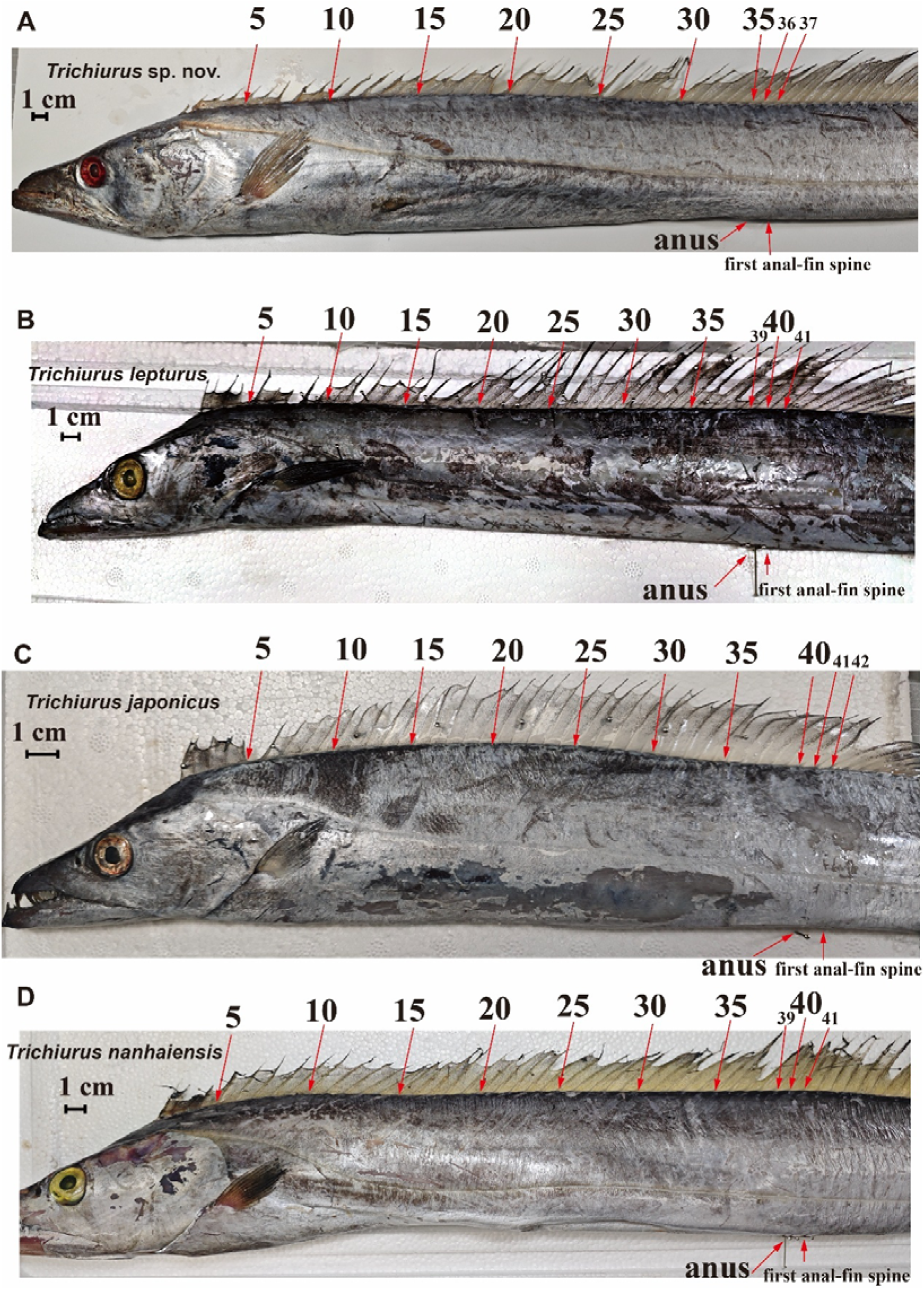
Position of anus and first anal-fin spine relative to the dorsal-fin rays in *Trichiurus* species. **A.** *Trichiurus javaensis* sp. nov., holotype (SOU18011027-15). **B.** *Trichiurus lepturus* (SOU18011028-2). **C.** *Trichiurus japonicus* (SOU18011029-9). **D.** *Trichiurus nanhaiensis* (market specimen, Shanwei City, Guangdong Province, China).

#### Paratypes

SOU18011027-1–SOU18011027-15 (15 specimens), 76.9–110.2 cm SL, 31.6–43.3 cm PL; collection data same as for holotype.

#### Diagnosis

*Trichiurus* species distinguished by following characters: dorsal-fin rays III, 116–125; anal-fin rays I, 95–105; first anal-fin pterygiophore situated below 36th–37th dorsal-fin ray; anus below 35th–36th dorsal-fin ray; anterior margin of pectoral-fin spine not serrated; fangs on both jaws with barbs; anteriormost tip of supraoccipital well posterior to posterior margin of the eye; no hyperostosis posterior to supraoccipital crest; upper jaw long (mean 16.6% of PL, range 15.5–17.6%); snout short (mean 12.0% of PL, range 10.9–13.1%); eye small (mean 5.3% of PL, range 4.3–5.7%). Pelvic and caudal fins absent.

#### Description

Morphometric and meristic data for type series of *T. javaensis* provided in Table 1. Body extremely elongated, ribbon-like, strongly compressed, tapering to a point posteriorly. Mouth large, tip of both the upper and lower jaws with a dermal process. Single nostril present on each side of head. Two pairs of enlarged, barbed fangs near tip of upper jaw; one pair near tip of lower jaw. Snout length distinctly less than upper jaw length. Interorbital space flat. Posteroventral margin of gill cover concave. Predorsal length slightly greater than body depth at dorsal-fin origin. Lateral line originates at upper margin of gill cover, runs obliquely to behind tip of pectoral fin, then descends to run parallel and close to the ventral body contour to the end of body. Distance between lateral line and ventral profile at anus slightly less than half distance to the dorsal profile. Pelvic and caudal fins absent.

**Table 1.** Comparative morphometrics and meristics of *Trichiurus javaensis* sp. nov. (holotype and paratypes) and related *Trichiurus* species.

|  | *T. javaensis* |  | *T. lepturus* | *T. nanhaiensis* | *T. japonicus* |
| --- | --- | --- | --- | --- | --- |
|  | Holotype | Paratype | SOU18011028- | SOU18011029-1~ | SOU18011029- |
|  | SOU1801 | SOU18011027- | 1~ | SOU18011029-6 | 7~ |
|  | 1027-16 | 1~SOU1801102 | SOU18011028- |  | SOU18011029- |
|  |  | 7-15 | 5 |  | 12 |
| Standard length(mm) | 1202.3 | 769.7-1102.0 | 704.8-852.3 | 616.7-816.5 | 648.9-840.2 |
| Head length (mm) | 157.2 | 98.4-139.5 | 107.2-114.5 | 77.4-107.6 | 68.2-95.0 |
| Preanal length (mm) | 472.2 | 316.2-433.9 | 310.1-329.0 | 240.4-322.7 | 201.4-276.1 |
| Dorsal fin rays | III, 125 | III, 116-125 | III, 131-136 | III, 132-135 | III, 131-138 |
| Anal fin rays | I, 105 | I, 95-105 | I, 101-105 | I, 98-105 | I, 98-113 |
| Pectoral fin rays | I, 11 | I, 11 | I, 10-11 | I, 10 | I, 10-11 |
| Dorsal fin rays opposite first anal spine | 36<sup>th</sup>-37<sup>th</sup> | 36<sup>th</sup>-37<sup>th</sup> | 40<sup>th</sup>-41<sup>st</sup> | 40<sup>th</sup>-41<sup>st</sup> | 41<sup>st</sup>-42<sup>nd</sup> |
| Dorsal fin rays opposite anus | 35<sup>th</sup>-36<sup>th</sup> | 35<sup>th</sup>-36<sup>th</sup> | 39<sup>th</sup>-40<sup>th</sup> | 39<sup>th</sup>-40<sup>th</sup> | 40<sup>th</sup>-41<sup>st</sup> |
| % in Preanal length (Mean) |  |  |  |  |  |
| Head length | 33.3 | 29.6-34.1 (31.6) | 33.5-34.9 (34.4) | 32.2-38.4 (35.8) | 33.9-38.2(36.1) |
| Upper jaw length | 17.6 | 15.5-17.3 (16.6) | 16.0-17.9 (17.2) | 17.0-18.3 (17.7) | 16.8-18.1(17.5) |
| Snout length | 11.4 | 10.9-13.1 (12.0) | 13.4-14.6 (14.1) | 12.9-14.2 (14.1) | 11.0-13.3(11.8) |
| Body depth at dorsal-fin origin | 15.4 | 15.2-19.2 (17.3) | 15.1-17.1 (15.8) | 15.9-18.3 (17.1) | 15.4-18.0(16.7) |
| Body width at dorsal-fin origin | 8.3 | 8.2-10.8 (9.2) | 7.1-8.3 (7.6) | 6.3-7.4 (6.8) | 5.8-7.9 (7.0) |
| Body depth at anus | 16.0 | 17.7-20.5(19.3) | 15.4-17.9 (17.2) | 18.1-20.3(19.4) | 18.2-19.9(18.8) |
| Body width at anus | 5.8 | 5.7-8.5 (7.0) | 4.8-5.9 (5.5) | 5.9-7.1(6.6) | 5.6-6.5(6.1) |
| Pectoral fin length | 12.1 | 10.7-14.8 (12.2) | 13.1-15.8 (14.2) | 11.8-14.0 (12.6) | 9.3-11.8 (11.1) |
| Pre-dorsal length | 22.0 | 19.1-21.4 (20.1) | 20.3-23.2 (21.9) | 21.4-23.8 (22.8) | 22.8-26.9(24.7) |
| Eye diameter | 4.3 | 5.0-5.7 (5.3) | 6.1-6.9 (6.4) | 5.6-6.2 (5.9) | 5.6-6.4 (6.0) |
| Interorbital width | 4.9 | 5.3-6.5 (5.8) | 4.4-5.9 (5.2) | 4.4-5.1 (4.8) | 4.4-5.2 (4.7) |

#### Color

Fresh specimens iridescent metallic on body, gradually fades to grayish-white after death. In preserved specimens, epidermis often detaches, revealing tan-brown subcutaneous layer. Dorsal fin semi-transparent white to tan-brown. Eyes white or red due to blood infiltration.

#### Distribution

*Trichiurus javaensis* currently known only from type locality: southern Indian Ocean, off coast of Yogyakarta, Indonesia.

#### Etymology

The specific epithet is derived from Java Island, the type locality of the new species.

#### Comparative Materials

SOU18011028-1–SOU18011028-5: *Trichiurus lepturus* Linnaeus, 1758; 5 specimens; Persian Gulf, coastal waters of Bandar Lengeh, Hormozgan Province, Iran (26°30’N, 55°00’E); 4 January 2025. SOU18011029-1–SOU18011029-6: *Trichiurus nanhaiensis* Wang & Xu, 1992; 6 specimens; Indian Ocean population, imported from Indonesia. SOU18011029-7–SOU18011029-12: *Trichiurus japonicus* Temminck & Schlegel, 1844; 6 specimens; East China Sea, near Haijiao Island, Zhoushan, Zhejiang Province, China (30°44’N, 123°09’E); 12 August 2025.

### Phylogenetic relationship

Phylogenetic trees were constructed based on mitochondrial *COX1* (Figure 3) and *16S rRNA* (Figure 4) sequences of all *Trichiurus* species available in the NCBI database that included geographic information for each specimen. Duplicate sequences were removed, only those representing monophyletic species or population groups were retained. The resulting phylogenetic relationships were used to distinguish *T. javaensis* from *T. lepturus*.

**Figure 3.**
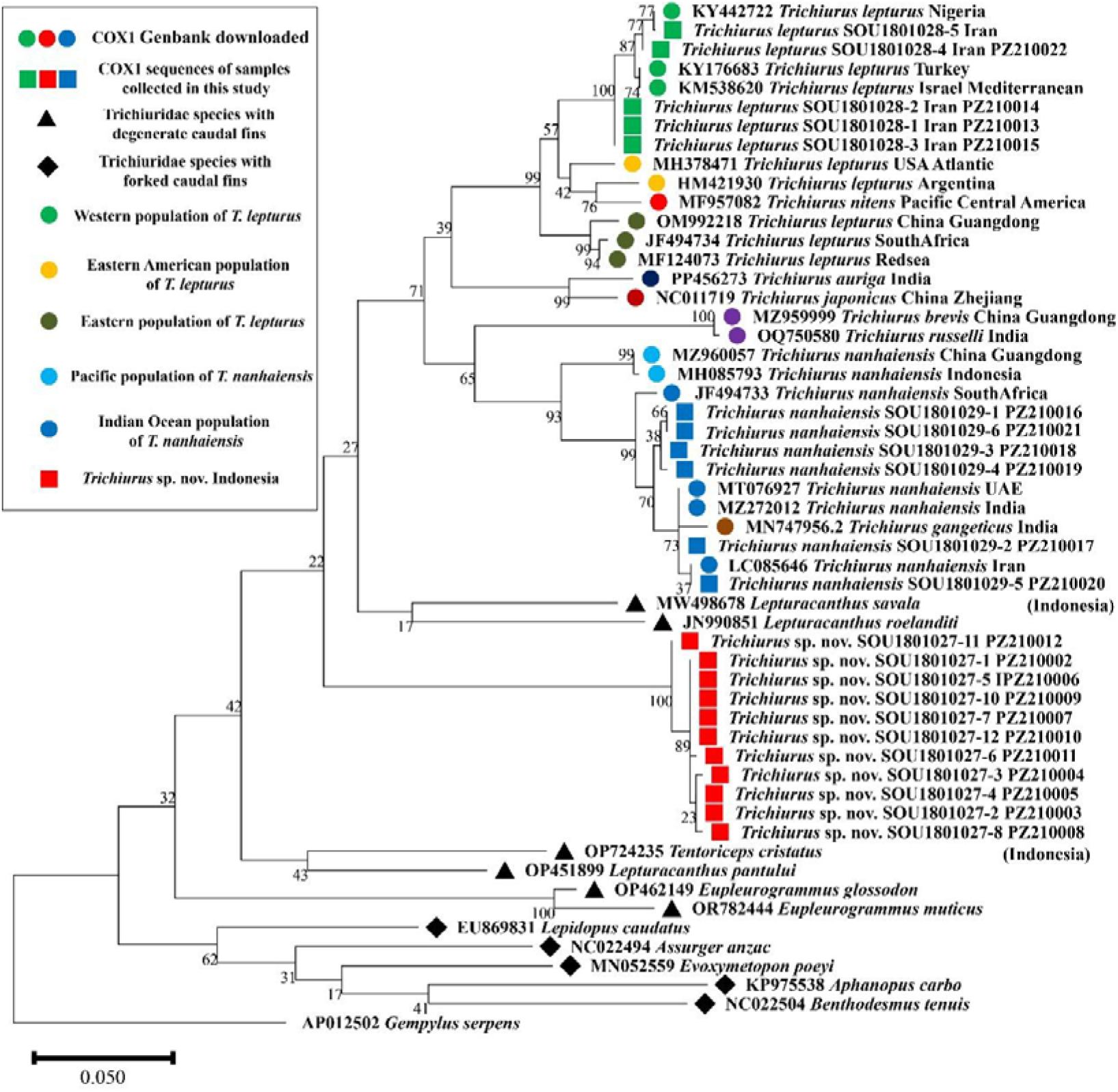
Maximum Likelihood phylogeny of family Trichiuridae based on mitochondrial COX1 gene sequences, with *Gempylus serpens* designated as outgroup. Numbers at nodes represent bootstrap support values from 1000 replicates. Bootstrap support values are expressed as percentages.

**Figure 4.**
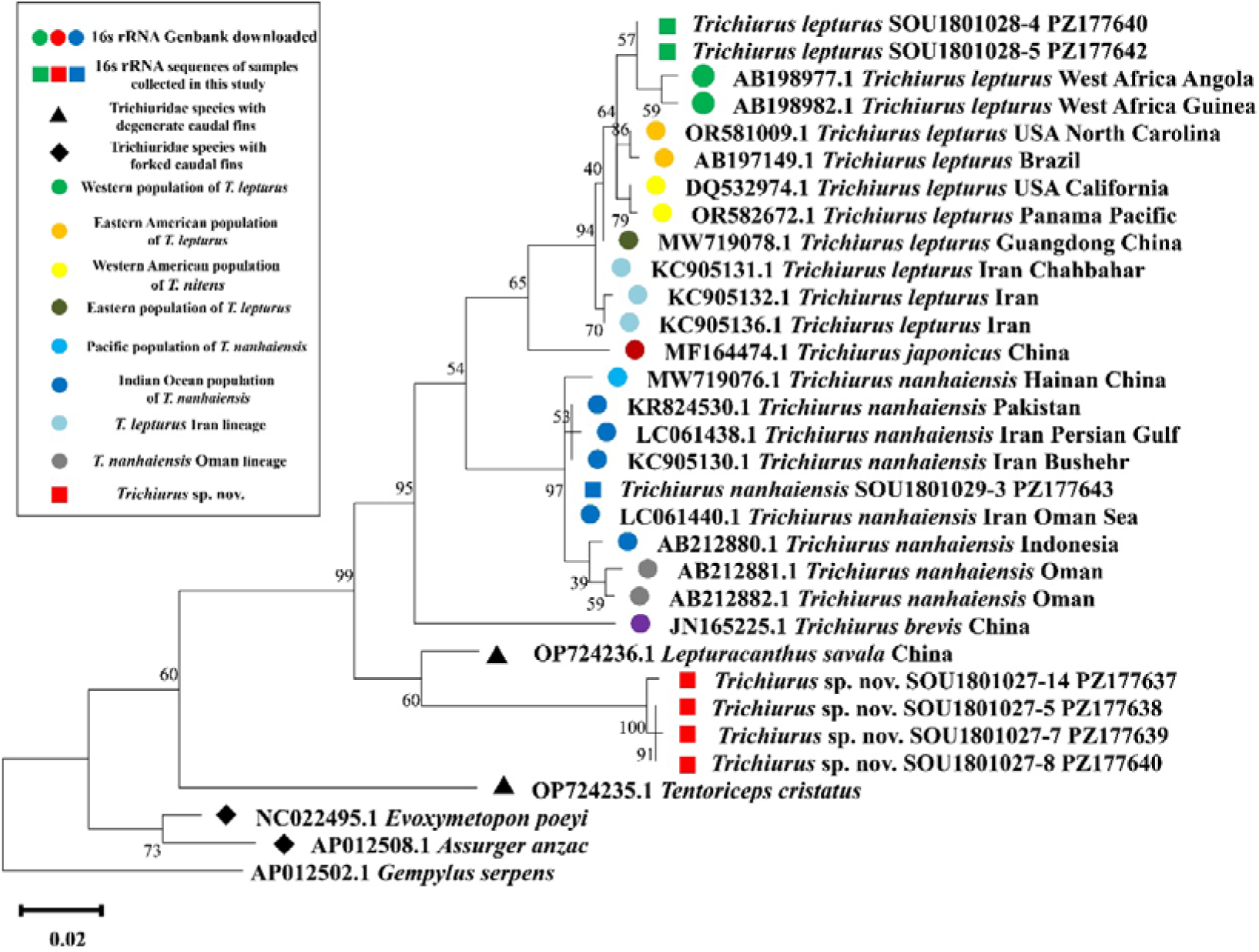
Maximum Likelihood phylogeny of family Trichiuridae based on mitochondrial 16S ribosomal RNA gene sequences, with *Gempylus serpens* the outgroup. Numbers at nodes represent bootstrap support values from 1000 replicates. Bootstrap support values are expressed as percentages.

In both phylogenetic trees, *T. javaensis* formed a distinct cluster outside the main *Trichiurus* clade and did not group with any other ribbonfish genera. Within *Trichiurus*, *T. japonicus*, *T. nanhaiensis*, and *T. brevis* each form distinct monophyletic clades, whereas specimens identified as *T. lepturus* cluster in the uppermost clade of the phylogeny. To visualize the geographic relationships of these lineages, sampling localities sharing monophyletic relationships are connected to produce a global distribution map of various “*T. lepturus*” lineages (Figure 5).

**Figure 5.**
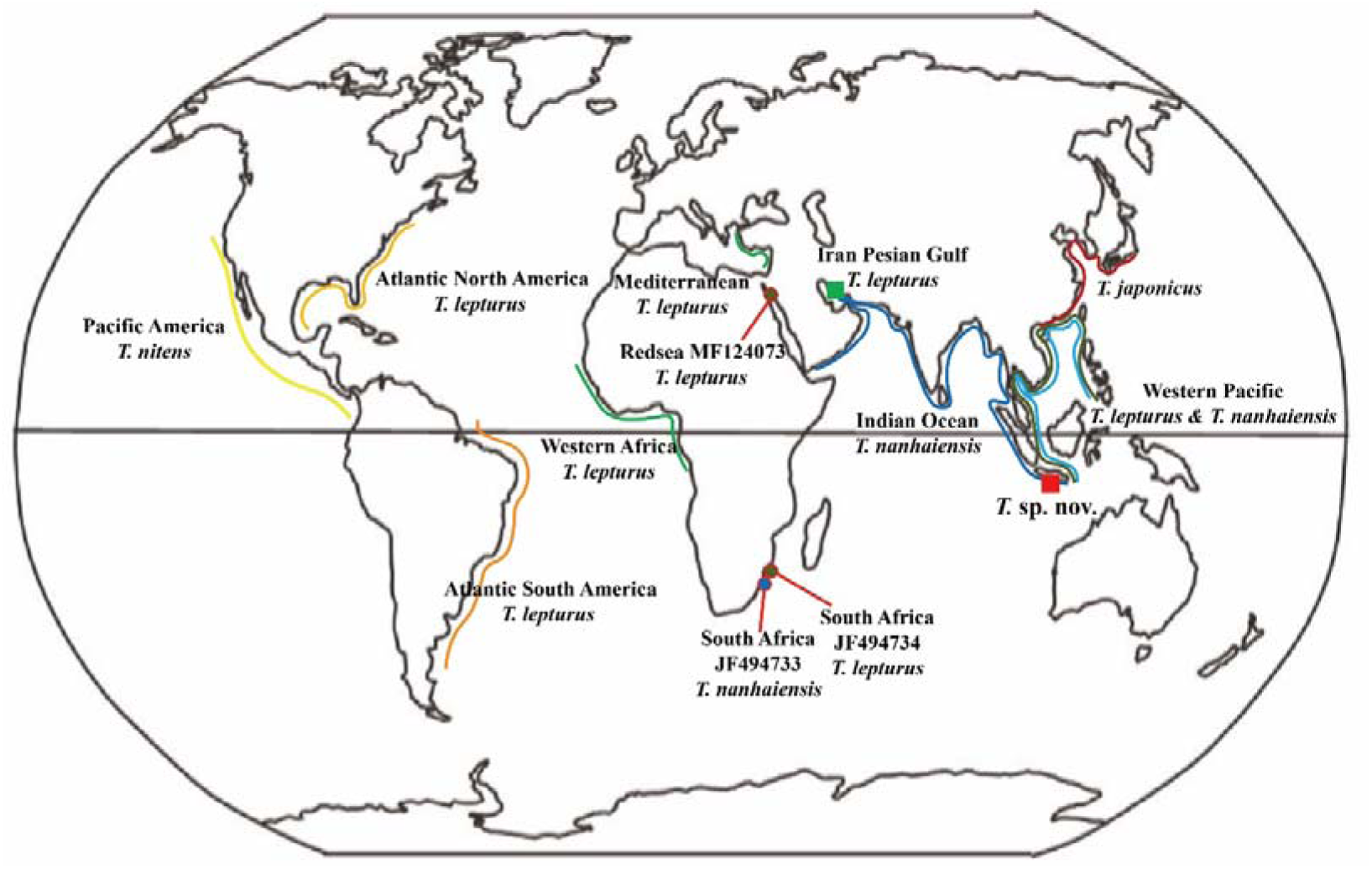
Global distribution of species and populations genetically identified as *Trichiurus lepturus*, based on 16S rRNA and COX1 sequence data.

The *T. nanhaiensis* lineage can be divided into two major groups: the western Pacific population (South China Sea, Vietnam, Thailand, and the Java Sea of Indonesia) and the Indian Ocean population (Yemen, Oman, Iran, Pakistan, India, Bangladesh, Myanmar, and Indonesia). Although these two groups show clear mitochondrial differentiation, their morphological differences are subtle. In addition, the *16S rRNA* sequence from Oman (AB212881; Chakraborty et al., 2006) and the *COX1* sequence from South Africa (JF494733) form two distinct monophyletic lineages, separate from both the western Pacific and Indian Ocean groups.

*Trichiurus lepturus* represents a globally distributed complex. Specimens from the Americas form three closely related monophyletic groups corresponding to the North Atlantic, South Atlantic (Brazil and Argentina), and Pacific (*T. nitens*) populations. Linnaeus’s type material from the United States and Jamaica most likely corresponds to the North Atlantic *T. lepturus*. Samples from the Mediterranean (Turkey, Israel), West Africa, and the Persian Gulf (Iranian specimens SOU18011028-1 to SOU18011028-5 from this study) form another western Old World clade, whereas those from the western Pacific (South China Sea, Philippines, Vietnam, and the Java Sea) cluster within a separate eastern clade. Additionally, *COX1* sequences from the Red Sea (Israel, MF124073) and South Africa (JF494733) form a lineage closely related to the western Pacific group. The *16S rRNA* sequence from the Gulf of Oman, Iran (KC905131) clusters in a previously unreported clade.

## Discussion

The newly described *T. javaensis* morphologically belongs to the *T. lepturus* complex, characterized by the highest point of the supraoccipital crest being positioned well behind the level of the posterior distal margin of the eye, a feature fundamentally distinguishes it from species of the *T. brevis* complex. Within the *T. lepturus* complex, *T. javaensis* can be differentiated by the position of the anus. In *T. lepturus*, *T. nanhaiensis*, and *T. japonicus*, the anus is situated below the 39th–41st dorsal-fin rays, whereas in *T. javaensis*, it is located below the 35th–36th dorsal-fin rays (Figure 2).

To establish *T. javaensis* as a distinct species, it is essential to compare it with other congeners, particularly *T. auriga* (Klunzinger, 1884), *T. gangeticus* (Gupta, 1966), and members of the *T. brevis* complex, which all possess well-defined diagnostic features. However, the critical comparison lies within the *T. lepturus* complex (*T. lepturus*, *T. japonicus*, *T. nanhaiensis*, and *T. nitens*).

Although *Trichiurus* species exhibit geographically structured diversity worldwide, populations from various regions have long been collectively referred to as *T. lepturus*. Based on Li’s (2006) re-examination of the four syntypes of *T. lepturus* described by Linnaeus in *Systema Naturae*, the single specimen preserved at the Swedish Museum of Natural History (NRM 15), designated as the lectotype, possesses fewer dorsal-fin rays (107) and a longer first anal-fin spine, indicating that it belongs to the genus *Lepturacanthus*. A second specimen, ZIU 172 from Uppsala University, was identified as a member of *Eupleurogrammus*, while the remaining two, originating from the United States and Jamaica, properly belong to *Trichiurus*. Linnaeus (1758) originally considered *T. lepturus* to be a cosmopolitan species ranging from the Americas to China, a view later followed by Tucker (1956), James (1967), and Nakamura & Parin (1993). This interpretation has led to the frequent misapplication of the name *T. lepturus* to three currently valid species: *T. japonicus* Temminck & Schlegel, 1844; *T. nanhaiensis* Wang & You, 1992; and *T. nitens* Garman, 1899.

*Trichiurus nanhaiensis* Wang & You, 1992 was described from the South China Sea as a valid species, differing morphologically from *T. japonicus* (referred to as *T. haumela* in their original description) by its yellowish dorsal fin and the bony hyperostosis on the head. *Trichiurus margarites* Li, 1992 was later synonymized with *T. nanhaiensis*. However, because Wang’s (1992) description was published in Chinese and based on limited material, *T. japonicus* and other regional forms of *T. lepturus* were not thoroughly compared, leading some subsequent studies to refer to this taxon as *T. sp. 2* or *T. lepturus nanhaiensis*.

*Trichiurus japonicus* Temminck & Schlegel, 1844, originally named *T. lepturus japonicus*, is distributed along the coasts of Japan, Korea, and China, and has one invalid synonym, *T. haumela*. Although mitochondrial data clearly separate *T. japonicus* from *T. lepturus*, no universally accepted morphological characters have yet been identified to distinguish them unambiguously.

*Trichiurus nitens* Garman, 1899, redescribed by Burhanuddin and Parin (2008), is a valid species distributed along the Pacific coasts of the Americas. It is characterized by the first anal-fin spine lying below the 36th–39th dorsal-fin rays. In *T. javaensis*, the anus is located below the 35th–36th dorsal-fin rays, and the first anal-fin spine lies below the 36th–37th dorsal-fin rays (Figure 2A, B), partially overlapping with *T. nitens*. However, *T. javaensis* differs by having a smaller eye diameter–to–preanal length ratio (5.1% [4.2–5.7%]) compared with *T. nitens* (7% [6–8%]), which does not overlap between the two species. Furthermore, *T. javaensis* exhibits a longer pre–upper-jaw length–to–anal length ratio (16.5% [15.3–17.6%]) than that of *T. nitens* (14% [12–14%]).

In this study, we integrated comprehensive morphological comparison and molecular phylogenetic analysis based on mitochondrial COX1 and 16S rRNA sequences, to formally described *Trichiurus javaensis* sp. nov., a previously unrecognized cryptic cutlassfish species from the southern coastal waters of Java Island, Indonesia, within the Coral Triangle global marine biodiversity hotspot. This new species is morphologically affiliated with the *T. lepturus* complex, and can be unambiguously distinguished from all other valid *Trichiurus* species by a unique suite of diagnostic traits, including the position of the anus and first anal-fin spine relative to dorsal-fin rays, proportional characteristics of snout, upper jaw and eye diameter to preanal length, as well as diagnostic skeletal and dentition features.

*T. javaensis* was placed within a lineage composed of genera with degenerated caudal fins rather than those with forked caudal fins, and suggesting deeper morphological and genetic divergence from other *Trichiurus* species. However, the morphological traits of this new species, namely the absence of a slit on the lower jaw and a short first anal-fin spine with length less than the pupil diameter, are diagnostic of the genus *Trichiurus* instead of *Lepturacanthus*. Although its monophyletic lineage is recovered outside the clade of *Lepturacanthus* in phylogenetic analyses, *Lepturacanthus* was originally erected as a subgenus within the genus *Trichiurus*. Accordingly, the placement of this species in the genus *Trichiurus* is justified. Meanwhile, this study systematically resolves the long-standing taxonomic confusion within the genus *Trichiurus*. We confirm that *T. lepturus*, long misrecognized as a single cosmopolitan species, actually constitutes a species complex comprising multiple geographically structured, genetically divergent lineages and valid species, and correct the historical erroneous relegation of numerous regional *Trichiurus* taxa as junior synonyms of *T. lepturus*. We further delineate the global phylogeographic pattern of the *T. lepturus* complex, and clarify the interspecific relationships and lineage differentiation within the genus *Trichiurus*.

This work fills the gap in the taxonomic research of *Trichiurus* in Indonesian archipelagic waters, and updates the global species diversity framework of the genus. The accurate species delimitation and robust phylogenetic framework established in this study provide an essential scientific basis for stock assessment, biodiversity conservation and region-specific fisheries management of Trichiuridae resources worldwide. Future investigations should prioritize under-sampled regions including the eastern African coast, Arabian Sea and Indonesian archipelagic waters, to uncover potential cryptic lineages and further resolve the systematic evolution of the genus *Trichiurus*.

## Conclusion

This study combined morphological comparison and mitochondrial COX1/16S rRNA phylogenetic analysis to formally describe *Trichiurus javaensis* sp. nov., a new cryptic cutlassfish from southern Java, Indonesia, with unique diagnostic traits separating it from all valid congeners. We resolved the long-standing taxonomic confusion in *Trichiurus*, confirming that the widely recognized cosmopolitan *T. lepturus* constitutes a species complex with geographically structured divergent lineages, and mapped its global phylogeographic pattern. This work fills a key research gap in Indo-Pacific *Trichiurus* taxonomy, providing a critical scientific basis for Trichiuridae fisheries management and biodiversity conservation.

## Data Availability

The datasets generated and analysed during the current study are available in the Dryad Digital Repository: https://doi.org/10.5061/dryad.XXXXXXXXX. GenBank accession numbers of COX1 sequences for this project are PZ210002–PZ210022, and those for 16S rRNA are PZ177637–PZ177643.

---

**Supplementary Table 1.**
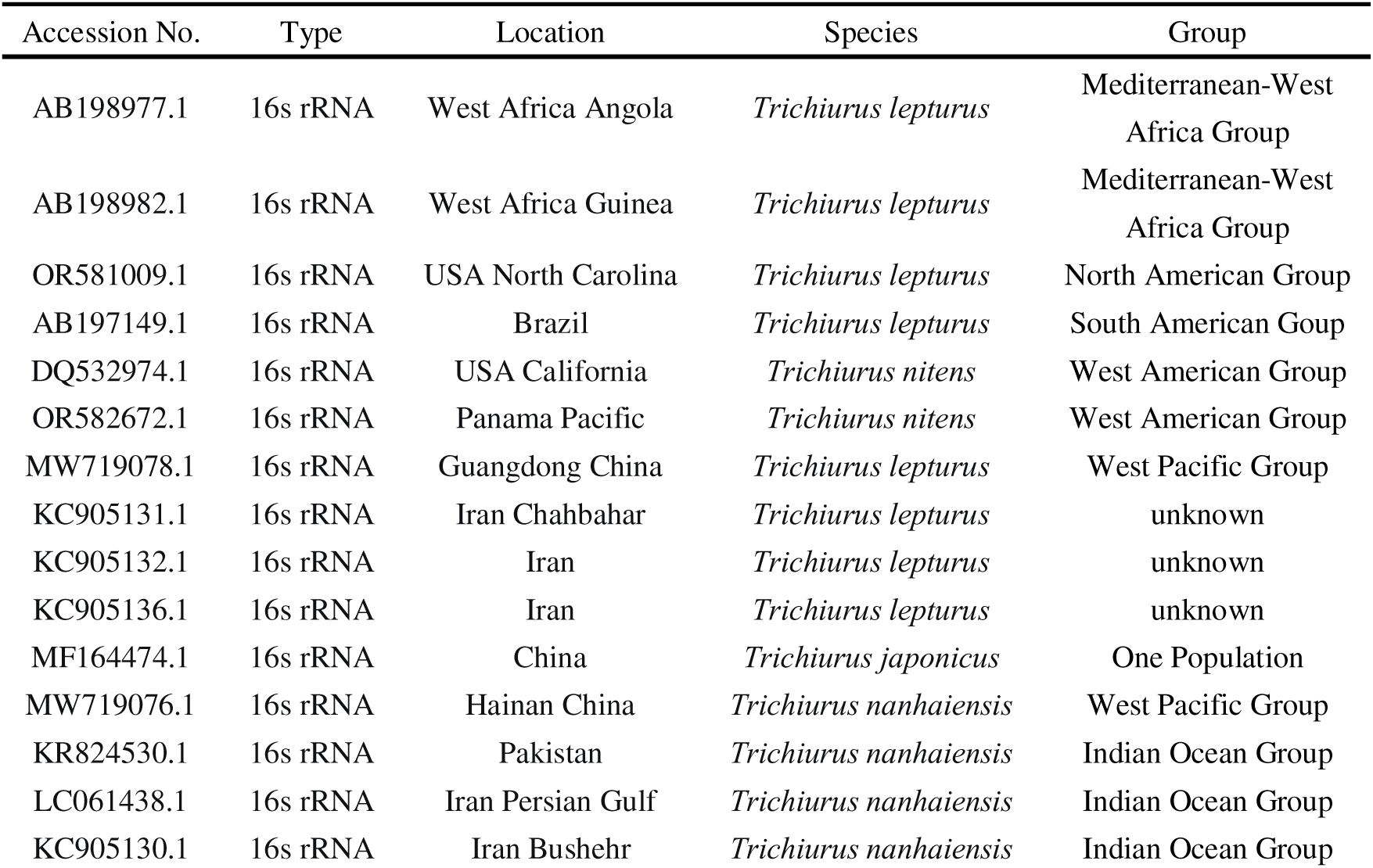
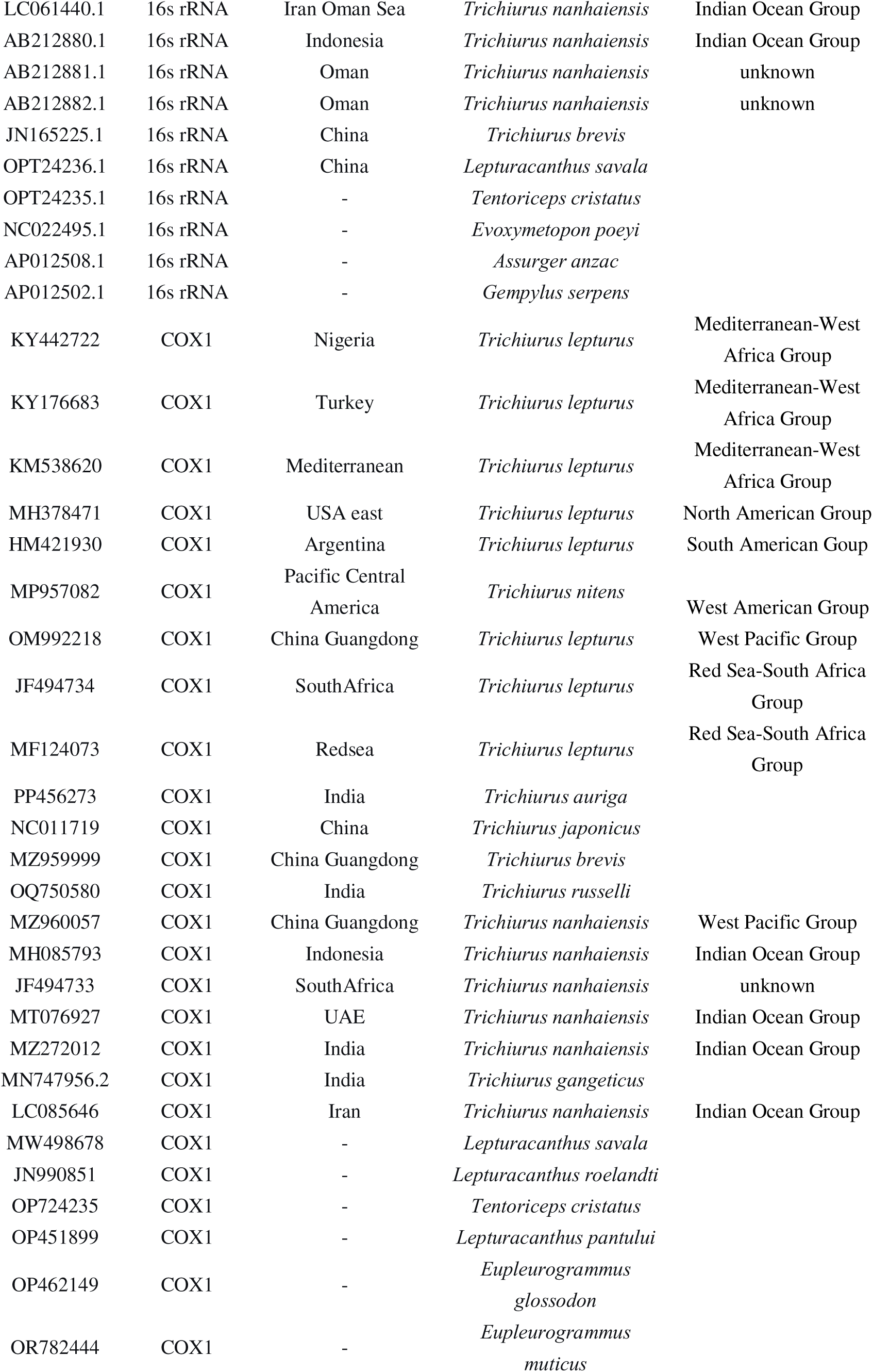
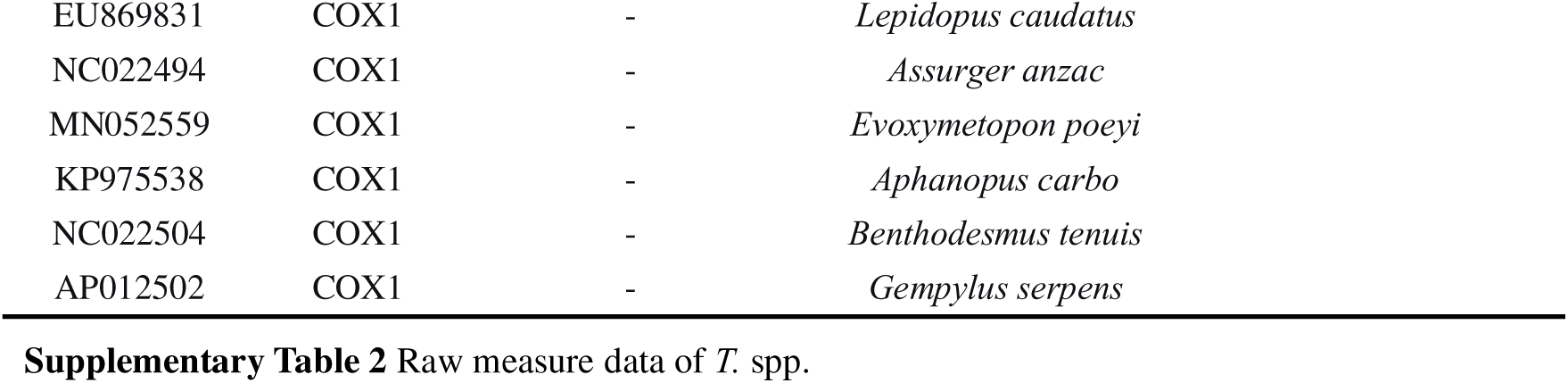
GenBank accession numbers and sample data for phylogenetic trees.

**Supplementary Table 2** Raw measure data of *T.* spp.

